# Structural basis of eIF2B-catalyzed GDP exchange and phosphoregulation by the integrated stress response

**DOI:** 10.1101/504654

**Authors:** Aditya A. Anand, Lillian R. Kenner, Henry C. Nguyen, Alexander G. Myasnikov, Carolin J. Klose, Lea A. McGeever, Jordan C. Tsai, Lakshmi E. Miller-Vedam, Peter Walter, Adam Frost

## Abstract

The integrated stress response (ISR) tunes the rate of protein synthesis. Control is exerted by phosphorylation of the general translation initiation factor eIF2. eIF2 is a GTPase, that becomes activated by eIF2B, a two-fold symmetric and heterodecameric complex that functions as eIF2’s dedicated nucleotide exchange factor. Phosphorylation converts eIF2 from a substrate into an inhibitor of eIF2B. We report cryoEM structures of eIF2 bound to eIF2B in the dephosphorylated state. The structures reveal that the eIF2B decamer is a static platform upon which one or two flexible eIF2 trimers bind and align with eIF2B’s bipartite catalytic centers to catalyze guanine nucleotide exchange. Phosphorylation refolds eIF2, allowing it to contact eIF2B at a different interface and, we surmise, thereby sequesters it into a non-productive complex.

**One Sentence Summary:** Structures of translation factors eIF2 and eIF2B reveal the mechanism of nucleotide exchange and its phosphoregulation during stress.

## Main Text

Initiation is the rate-determining step for translation of the genetic code into proteins, the macromolecules that perform catalytic, structural, and signaling functions in cells. Translation takes place at the ribosome, a macromolecular machine that utilizes amino acid-charged tRNA molecules that decode mRNA and catalyze sequential peptide bond formation. Numerous factors aid in this process, one of which is eukaryotic translation initiation factor 2 (eIF2), a heterotrimeric GTPase composed of α, β, and γ subunits. During translation initiation, eIF2 binds methionine initiator tRNA and GTP to form a ternary complex that scans mRNAs for start codons. Following start codon detection, eIF2γ hydrolyzes its GTP, translation begins, and eIF2-GDP dissociates from the ribosome. GDP must be released before eIF2 can initiate another round of translation and, since GDP release is intrinsically slow, the dedicated guanine nucleotide exchange factor (GEF), eIF2B, catalyzes this reaction to enable new protein synthesis.

Together, eIF2 and eIF2B form a regulatory control node. Stress-responsive kinases phosphorylate eIF2α at conserved serine 51 (or serine 52 in humans, if the leading Met residue is retained). This post-translation modification transforms eIF2 from a substrate into a competitive inhibitor of its own GEF, eIF2B, precluding the reloading of eIF2 with GTP and methionine-charged tRNA. This phosphoregulation of eIF2 is known as the Integrated Stress Response (ISR) (*1*). Once activated, the ISR reduces overall protein synthesis, while concomitantly enhancing the translation of a small subset of mRNAs, such as ATF4, in response to cellular threats like protein misfolding, infection, inflammation, nutrient deprivation, and other inputs (*1*–*3*).

eIF2B is a remarkably complex nucleotide exchange factor. The holoenzyme comprises two copies each of an α, β, γ, δ, and ɛ subunit that assemble into a two-fold symmetric heterodecamer (*4*, *5*). The eIF2Bε subunit contains the enzyme’s catalytic center and associates most closely with eIF2Bγ. Two copies each of the structurally homologous eIF2Bβ and δ subunits directly interface within the core, while a homodimer of two eIF2Bα subunits bridges across the symmetry interface of the decamer (*5*, *6*). Genetic and biochemical studies have identified conserved residues responsible for eIF2B’s catalytic activity and have suggested how eIF2 binding to eIF2B may differ following eIF2α-S51 phosphorylation (*5*, *7*–*11*). However, many questions remain outstanding, including the mechanistic basis of eIF2 recognition by eIF2B and eIF2B’s mechanism of catalyzing guanine nucleotide exchange. Moreover, the transformation of eIF2 from a substrate to high-affinity inhibitor of eIF2B following phosphorylation has remained enigmatic, despite its central importance to the ISR.

A small molecule inhibitor of the integrated stress response, ISRIB, was found to allay the effects of eIF2α phosphorylation by activating eIF2B (*12*–*14*). Upon adding ISRIB, translation is restored in cells undergoing the ISR, and translation of stress-responsive transcripts like ATF4 is repressed (*13*, *14*). When administered to rodents, ISRIB enhances cognition and reverses cognitive deficits due to traumatic brain injury (*15*) and prion-induced neurodegeneration (*16*). Furthermore, eIF2B activation rescues cognitive and motor function in mouse models of leukoencephalopathy with vanishing white matter (VWM), a debilitating or fatal autosomal disorder associated with hundreds of mutations in all subunits of eIF2B (*17*, *18*). Remarkably, ISRIB exerts its effects without overt toxicity (*12*, *16*) due to restoration of the cytoprotective activity of the ISR when stress levels increase (*19*).

Structural data show that ISRIB acts as a molecular staple, bridging the symmetric interface of two eIF2B subcomplexes to enhance the formation of the decameric eIF2B holoenzyme (*20*, *21*). Biochemical experiments showed that ISRIB enhances incorporation of the eIF2Bα subunit into the decamer but does not make direct contacts with this subunit (*20*). Therefore, we proposed that ISRIB’s enhancement of GEF activity derives from its ability to promote higher-order assembly of the eIF2B decamer—but without a structural understanding of how eIF2 binds to its GEF, it has remained a mystery why the decameric eIF2B would be more active than its unassembled subcomplexes. In addition, a structure of eIF2B’s inhibited state during the ISR could explain ISRIB’s bell-shaped response profile to increasing stress (*19*). To explore the relationship between eIF2B and eIF2, we embarked upon a structural study of eIF2B in combination with both its substrate and phosphorylated inhibitor.

### eIF2B heterodecamer bound to one or two eIF2 heterotrimers

We co-expressed all five subunits of human eIF2B in *Escherichia coli* and all three subunits of human eIF2 in *Saccharomyces cerevisiae* (Fig. S1A-B). The yeast expression strain lacked the only yeast eIF2 kinase (*gcn2Δ*) to ensure homogenous purification of the non-phosphorylated form of eIF2 (*22*). Given that ISRIB-stabilized eIF2B decamers contain two putative sites for nucleotide exchange, we sought structures that could illuminate the enzyme’s potential for cooperativity. We incubated ISRIB and the purified complexes together at concentrations near the Michaelis constant of the nucleotide exchange reaction (K_m_ = 1.5 µM, (*20*)) and then added the inter-amine crosslinker BS3 to stabilize potentially transient interactions before sample vitrification. As a result of this approach, cryoEM imaging, image classification, and reconstruction revealed two distinct structures. First, 44,157 particles contributed to an asymmetric reconstruction of eIF2B bound to a single eIF2 trimer (Fig. 1A-C, Figs. S2-S3, Table S1-S3). Second, 7,473 particles contributed to a reconstruction of eIF2B bound to two symmetrically arranged eIF2 trimers (Fig. 1D-F, Figs. S2-S3, Table S1-S3). Clear density for ISRIB indicated that the small molecule retained its symmetric orientation and binding site interactions as previously published (*20*, *21*). ISRIB’s binding pocket retained two-fold symmetry, unperturbed by asymmetrically bound eIF2 (Fig.1A, D).

**Fig. 1.**
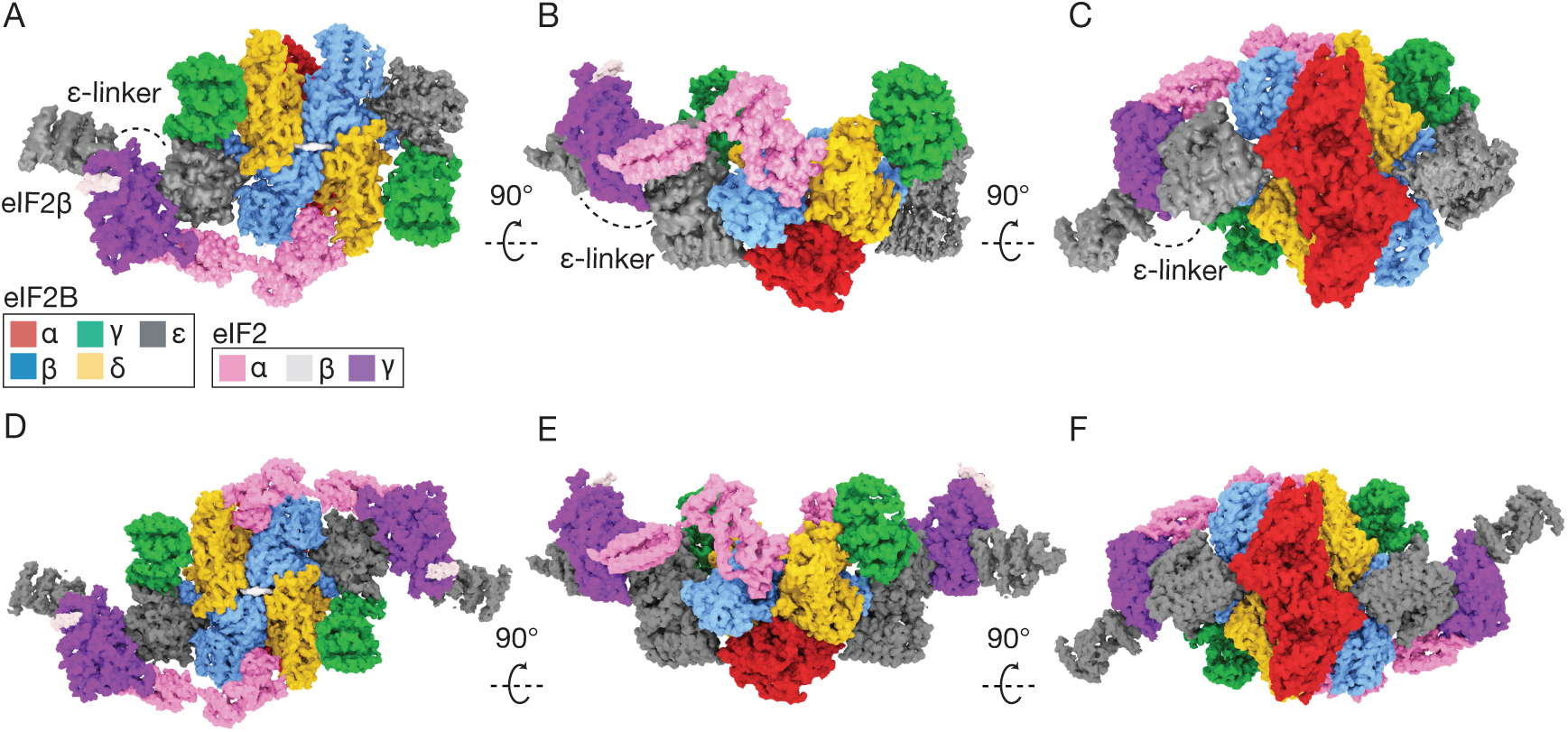
eIF2B heterodecamer bound to one or two eIF2 heterotrimers. (A-C) Orthogonal views of a single versus (D-E) a pair of elongated eIF2 heterotrimers bound to ISRIB-stabilized eIF2B decamers. ISRIB density is rendered in white.

Snaking across the surface of eIF2B, we observed cryoEM density consistent with the size and shape of eIF2 subunits as well as the previously unresolved HEAT domain of eIF2Bɛ.

Comparison with crystal and NMR structures of yeast and human eIF2α (PDBs: 1Q46,1Q8K) (*23*, *24*), and crystal structures of the yeast and human eIF2Bɛ HEAT domain (PDBs:1PAQ, 3JUI) (*25*, *26*), revealed that the fully assembled eIF2•eIF2B complex retained similarity to the structures of these independently purified domains. We only identified a single helix of eIF2β, indicating that the majority of the β subunit remained conformationally unconstrained in these assemblies (Fig. 1A, D), consistent with other studies (*27*, *28*). Finally, the more distantly related structure of the *Sulfolobus solfataricus* eIF2γ homolog, aIF2γ (PDB: 4RCY, 4RD6) (*29*), displayed remarkable similarity to the cryoEM density of the human subunit (Fig. 1-2).

In both of these reconstructions, all five subunits of eIF2B can be superimposed on previously solved structures determined in the absence of eIF2 (RMSD of ~0.6Å) (*20*). Also consistent with previous reconstructions of eIF2B, we resolved density for the “ear” domains of the eIF2Bγ subunits, but at a diminished resolution that precluded confident interpretation. eIF2 retained its overall structure and arrangement in the single- and the double-bound reconstructions (Fig. 1A-C versus D-F). These observations indicate that eIF2 binds via equivalent modes to both sides of a static eIF2B scaffold. The lack of conformational allostery in eIF2B upon eIF2 engagement is consistent with the non-cooperative kinetics reported for nucleotide exchange by fully assembled eIF2B decamers (*20*).

### Bipartite basis of guanine nucleotide exchange by eIF2B

When bound to eIF2B, the eIF2 heterotrimer adopted an elongated conformation (Fig. 2A) when compared to its tightly-wrapped arrangement around Met-tRNAi (*30*). The eIF2 subunits are stretched out across a 150Å span of eIF2B (Fig. 1-2). eIF2’s central nucleotide-binding γ-subunit is flanked by its α- and β-subunits at its opposing ends. eIF2γ contains motifs found in other GTP-binding proteins, including the nucleobase binding G4 motif, the P loop (phosphate binding), and switch helices 1 and 2. By contrast with these commonalities between eIF2 and other G-proteins, eIF2B has unique features consistent with the extraordinary diversity found in other GEF•GTPase structures (*31*). eIF2B recognizes eIF2 via coincident binding of both eIF2α and eIF2γ, and binding to each of these subunits involves bipartite elements of eIF2B, as we will discuss in turn (Fig. 1, Fig. 2A).

**Fig. 2.**
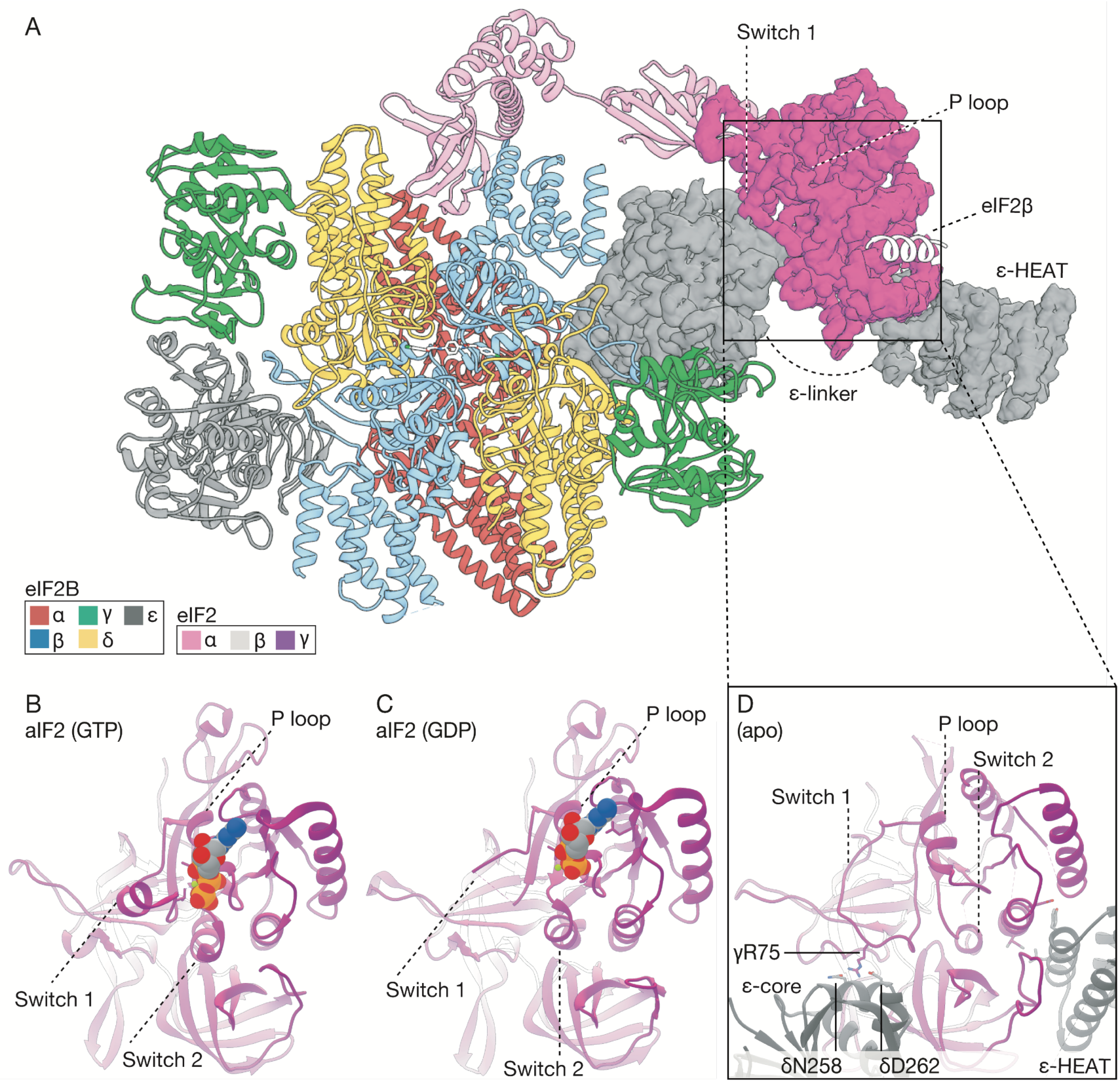
The bipartite basis of guanine nucleotide exchange by eIF2B. (A) Structural model of a single eIF2 heterotrimer bound to the eIF2B decamer, emphasizing the cryoEM density for eIF2γ and its interactions with the eIF2Bɛ subunit. (B) Comparison of an aIF2 structure bound to GTP (PDB: 4RCY), or (C) GDP (PDB: 4RD6), with the open and nucleotide-free state of eIF2 (D) reported here.

In the first example of bipartite recognition revealed by these structures, eIF2γ binds to two domains of eIF2Bɛ—separated by a flexible tether—which appear to function together to splay open the nucleotide binding site. Alignment of our nucleotide-free cryoEM model and the γ-subunit of GTP- and GDP-bound aIF2 from *Sulfolobus solfataricus* (*29*) revealed overall structural conservation (Fig. 2B-D, average RMSD of ~2.3Å). Within the confines of the GTP/GDP binding pocket, however, the structures diverged considerably (RMSD of ~12Å). Prior work has implicated the HEAT domain in catalysis (*25*, *26*). Consistent with these reports, we observed direct interactions between eIF2γ and the HEAT domain, especially with a partially hydrophobic surface that includes eIF2Bɛ Y583 and A584. These contacts appeared to “pull” a pair of α helices in eIF2γ, including the switch 2 helix, towards the HEAT domain and away from the nucleotide binding site (Fig. 2B-D). eIF2’s GTPase activating protein eIF5 contains a structurally homologous HEAT domain (*32*, *33*), suggesting that similar contacts may come into play during GTP hydrolysis as eIF2 progresses through translation initiation.

On the opposing side of the nucleotide binding pocket, we found the central core of eIF2Bɛ directly engaged with the switch 1 element of eIF2γ, which was rearranged into a well-resolved loop. Among the clearest interactions responsible for this change appear to be electrostatic interactions between eIF2γ R75 in switch 1 and eIF2Bɛ residues N258 and D262. Thus, both the HEAT domain and the core of eIF2Bɛ work together to pull the nucleotide binding site open, contributing to overall GEF activity (Fig. 2B-D).

### Bipartite basis of eIF2α recognition and assembly-stimulated activity

The second example of bipartite recognition revealed by these structures concerns eIF2α binding in the cleft between the eIF2Bβ and eIF2Bδ’ subunits (δ’ to indicate the δ-subunit from the opposing tetramer, Figs. 1-3). Notably, this bipartite binding site only exists after two tetramers of eIF2B(βγδɛ) associate to form the symmetry interface in the octamer eIF2B(βγδɛ)_2_.

eIF2α contains two structured domains separated by a flexible linker. The interaction with eIF2B just described occurs via its N-terminal domain, which consists of a five-stranded β-barrel oligonucleotide binding motif (OB-fold) (*23*), common to other tRNA binding proteins. The OB-fold is further elaborated with an α-helical subdomain connected by a conserved positively-charged loop (the S-loop), while the C-terminal αβ-fold domain connects eIF2α to eIF2γ. The S-loop harbors S51 and is responsible for all of the resolvable contacts between eIF2α and the eIF2B β subunit (Fig. 3A). For the *trans-*tetramer interaction, eIF2α exploits a conserved KGYID motif previously shown to be important for binding (*11*). Of note, an interaction between Y81 of this motif was well-resolved adjacent to the equally prominent R250 on eIF2Bδ’ (Fig. 3B). At a distance of ~4.3Å, this interaction constitutes a cation-π bond (Fig. 3A-B). To investigate the importance of this bond, we mutated R250 to either alanine or glutamate. Neither mutation affected the residual GEF activity displayed by dissociated tetramers (Fig. 3D, R250A k_obs_ = 0.013 min^−1^, R250E k_obs_ = 0.023 min^−1^, wild-type k_obs_ = 0.016 min^−1^), while both mutants diminished the GEF activity of the ISRIB-stabilized eIF2B octamer when compared to wild-type (Fig. 3E, R250A k_obs_ = 0.012 min^−1^, R250E k_obs_ = 0.017 min^−1^, wild-type k_obs_ = 0.063 min^−1^). These results are consistent with the notion that eIF2α interactions with the *trans*-tetramer only occur upon assembly of eIF2B’s symmetry interface in the octamer.

**Fig. 3.**
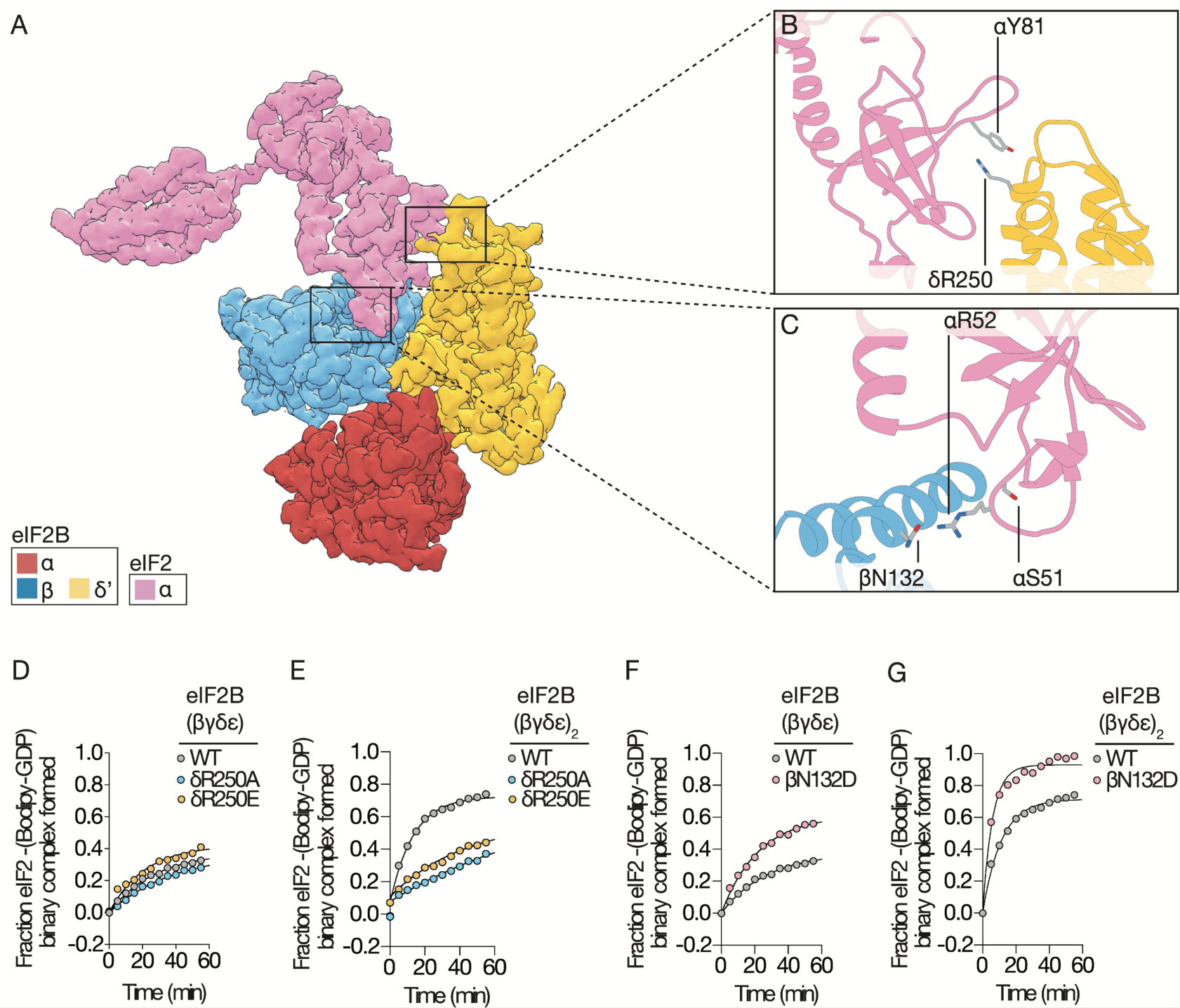
The bipartite basis of eIF2α recognition and assembly-stimulated activity. (A) cryoEM density for eIF2α bound to the regulatory subcomplex (α,β,δ or RSC) of eIF2B. (B) Zoom-in on a cation-π interaction between eIF2Bδ and eIF2α, and (C) polar interactions between eIF2Bβ and the S-loop of eIF2α. (D,F) GEF activity of wildtype versus mutated eIF2B(βγδε) tetramers, or (E,G) ISRIB-stabilized eIF2B(βγδε)_2_ octamers measured by loading of fluorescent GDP onto eIF2.

On the *cis*-tetramer, eIF2α’s positively-charged S-loop binds negatively charged and polar residues along the exposed surface of eIF2Bβ. This binding site is consistent with existing biochemical and genetic data from yeast suggesting that mutations in this site compromise binding to eIF2 (*7*). Examination of the structure identified a salt bridge between eIF2Bβ E135 and eIF2α R53, and a potential hydrogen bond between eIF2Bβ N132 and eIF2α R52 (Fig. 3C). To further explore the functional importance of the latter interaction, we substituted N132 with an aspartate, anticipating that the introduced charge complementarity would further enhance the interaction with eIF2α R52. When compared to wild-type eIF2B tetramers, eIF2B-βN132D tetramers and ISRIB-stabilized octamers proved to be gain-of-function mutations that exhibited two-fold enhanced GEF activity, consistent with an enhanced substrate interaction (Fig. 3F-G, Fig. S1D, eIF2B(βγδɛ) βN132D k_obs_ = 0.044 min-^1^, eIF2B(βγδɛ)_2_ βN132D k_obs_ = 0.169 min-^1^).

By intermolecular bridging, eIF2 thermodynamically stabilizes the eIF2B octamer. This structural observation is consistent with positive cooperativity in substrate binding (*5*) because the binding of the first eIF2 will create a symmetry-equivalent binding site on the other side of the octamer (Figs. 1 and 3). This observation is also consistent with eIF2B tetramers possessing significantly reduced catalytic activity when compared to the fully assembled decamer. Finally, eIF2α binding in the cleft between tetramers supports our proposal that ISRIB enhances eIF2B’s GEF activity by promoting the higher-order assembly of eIF2B.

By contrast with previous reports (*5*), the eIF2α N-terminal domain does not contact eIF2Bα in our structure, implying that a major role for eIF2Bα is to facilitate nucleotide exchange using a mechanism that is analogous and synergistic with ISRIB’s activity: favoring the association of eIF2B(βγδɛ) tetramers into octamers (upon ISRIB binding) and decamers (upon eIF2Bα_2_ binding), but not through direct substrate interactions. We surmise that by this mechanism ISRIB and/or eIF2Bα_2_ binding enhance nucleotide exchange by positioning the GDP-bound eIF2γ subunit in proximity to one of the two catalytic domains of eIF2Bɛ available in the complex. With dissociated tetramers, only one side of the eIF2α binding site remains, reducing binding and, in turn, limiting GEF activity. Together these data point to the regulated assembly of eIF2B decamers as a critical node for tuning eIF2B GEF activity in response to cellular needs.

### Structural basis of phosphoregulation by the ISR

To understand how eIF2α phosphorylation on S51 transforms eIF2 from a substrate into a competitive inhibitor of eIF2B, we co-expressed and purified the isolated eIF2 α subunit in *Escherichia coli* together with the kinase domain of PERK, one of the four ISR-inducing kinases (Fig. S1C). Phospho-specific Western blotting and Phos-tag gel electrophoresis confirmed complete and specific phosphorylation on S51 (Fig. S1C). We incubated pre-assembled eIF2B decamers with a 3-fold molar excess of eIF2α-P, crosslinked with BS3, and vitrified for cryoEM imaging. After classification and refinement, 22,165 particles contributed to a 2-fold symmetric reconstruction of the eIF2B decamer adorned with a two copies of eIF2α-P (Fig. 4A, Figs. S2-S3, Tables S1-S3). In both cases, eIF2α-P bridges the interface between eIF2Bδ and eIF2Bα (Fig. 4A). Intriguingly, we observed no overlap between the binding sites of non-phosphorylated eIF2α observed in the eIF2•eIF2B structure described above (Fig. 1-3) and eIF2α-P (Fig. 4B-C).

**Fig. 4.**
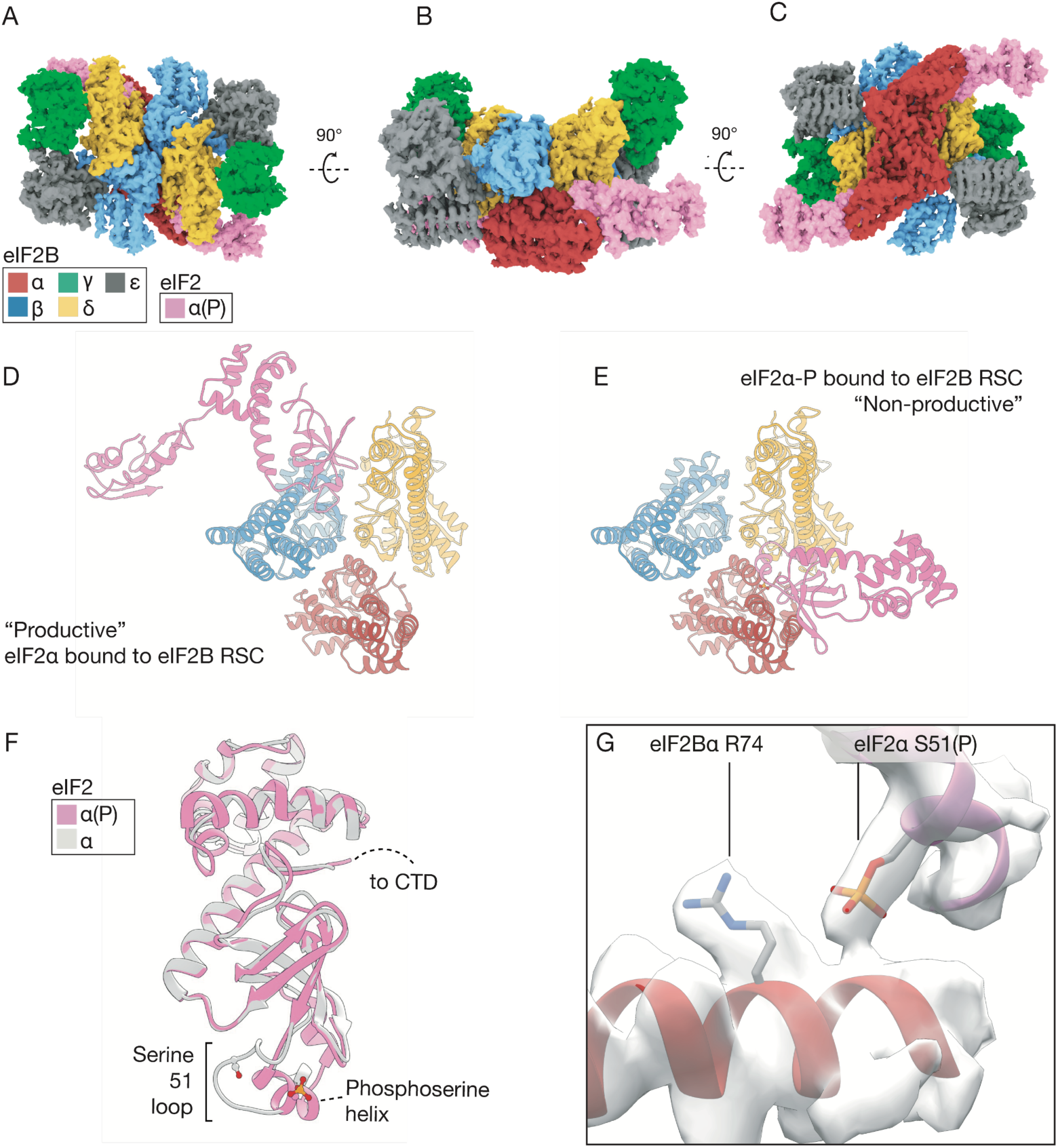
The structural basis of phosphoregulation by the ISR. (A-C) Orthogonal views of a pair of S51-phosphorylated eIF2α subunits bound to the eIF2B decamer. (D) Comparison of the productive binding mode of non-phosphorylated eIF2α, versus (E) the non-productive and non-overlapping binding mode of phosphorylated eIF2α. (F) Overlay comparison of the refolded conformation of the S-loop in the non-versus the phosphorylated states. (G) CryoEM density and interpretation, placing the S51 phosphate moiety near eIF2Bα R74.

At ~3.3Å resolution, density for both eIF2α S51-P and an arginine 5.7Å away, eIF2Bα R74, were clearly resolved and indicative of a long-range electrostatic interaction (Fig. 4G).

Moreover, phosphorylation of S51 induced refolding of the S-loop into a short α-helix (Movie S1, Fig. 4F). The phosphorylation-induced conformational rearrangement positions hydrophobic residues on eIF2α for direct interactions with surface-exposed hydrophobic residues on eIF2Bα and eIF2Bδ (including eIF2α I55, I58, and L62, Movie S1).

This new binding mode is consistent with biochemical analyses and numerous known mutations from *in vivo* yeast and mammalian studies. First, eIF2Bα is dispensable for viability in yeast, but deletion of this gene impairs phospho-inhibition of eIF2B, consistent with the subunit’s role in binding eIF2α-P in our structure (*34*). Point mutations with identical phenotypes cluster at the interface between eIF2Bα and eIF2Bδ, for example eIF2Bα F239 and eIF2Bδ M506 and P508 (*35*, *36*). Importantly, eIF2Bδ L314 complements the hydrophobic surface of the eIF2α S-loop that is exposed upon refolding, and mutation of the equivalent position in *S. cerevisiae*, L381Q, impairs the ISR in yeast (*35*). Finally, eIF2Bα T41, which also contributes to the eIF2α-P binding surface, is critical for the maintenance of the ISR in both yeast and mammalian studies (*34*). Together these data validate the phosphorylation-induced refolding and relocation of eIF2α-P observed in our structure that corresponds with the conversion of eIF2 from substrate to inhibitor of eIF2B-catalyzed nucleotide exchange.

## Discussion

Our analyses reveal the mechanistic basis of eIF2B’s nucleotide exchange activity and suggest how phosphorylation converts eIF2 from a substrate into an inhibitor. eIF2α subunit recognition in the cleft between adjacent tetramers of eIF2B(βγδɛ) productively positions the eIF2γ subunit so that it can be pulled open by eIF2Bɛ, stabilizing the nucleotide-free state. Since the concentration of GTP in the cell is significantly higher than that of GDP, after release of eIF2 from eIF2B, GTP will preferentially bind to the empty site, allowing eIF2 to engage in another round of ternary complex formation. Prior crosslinking studies suggested that eIF2 binds across the decameric interface, engaging the eIF2α subunit with eIF2Bβ and δ subunits from opposing tetramers (8), and binding studies reflect the importance of this assembly event (*37*). Here we show that the non-phosphorylated form of eIF2 binds to a composite surface spread across the assembled decamer and that this allows both the core and the flexibly attached HEAT domain of eIF2Bɛ to engage its target in concert for enhanced GEF activity.

By contrast, S51-phosphorylated eIF2α-P adopts a new helical conformation such that the S-loop becomes incompatible for binding to the site where nonphosphorylated eIF2α binds as a substrate. Rather, phosphorylation enables an entirely distinct binding mode on the opposite side of eIF2B where eIF2α-P lies exiled at the interface of eIF2Bα and eIF2Bδ. This new binding site depends on an electrostatic interaction between eIF2Bα R74 and the phosphate moiety as well as a conserved hydrophobic surface that is exposed upon phosphorylation-induced refolding. We surmise that this new binding mode is nonproductive for nucleotide exchange on eIF2-P, and also sequesters the GEF in an inhibited state, perhaps by occluding one or both of the two critical regions of eIF2Bɛ.

According to this view, the decameric core of eIF2B is a static scaffold. Control is exerted by regulation of its assembly state, which can be influenced by ISRIB and eIF2Bα_2_. Additional control arises from binding of eIF2-P which binds as a competitive inhibitor but to a non-overlapping binding site for the alpha subunit. We note that eIF2-P inhibitor binding requires the presence of eIF2Bα_2_ with which it interacts directly, whereas eIF2 substrate binding does not, as it can occur with the octamer. Atomic models of ISRIB-stabilized eIF2B bound to eIF2 reconcile numerous structure-activity relationships and are consistent with both loss- and gain-of-function mutations described here and previously. Together, the structures provide an intuitive view of how holoenzyme assembly activates nucleotide exchange as well as provides opportunities for regulation (Fig. 5). Both the substrate eIF2α subunit, the regulatory eIF2Bα_2_ subunit, and ISRIB stabilize the two-fold symmetric and fully active decameric form of eIF2B by “stapling” the constituents together. By bridging between eIF2Bα and eIF2Bδ at its binding site, eIF2-P is likewise predicted to stabilize the eIF2B decamer, albeit now holding it in an inactive state.

**Fig. 5.**
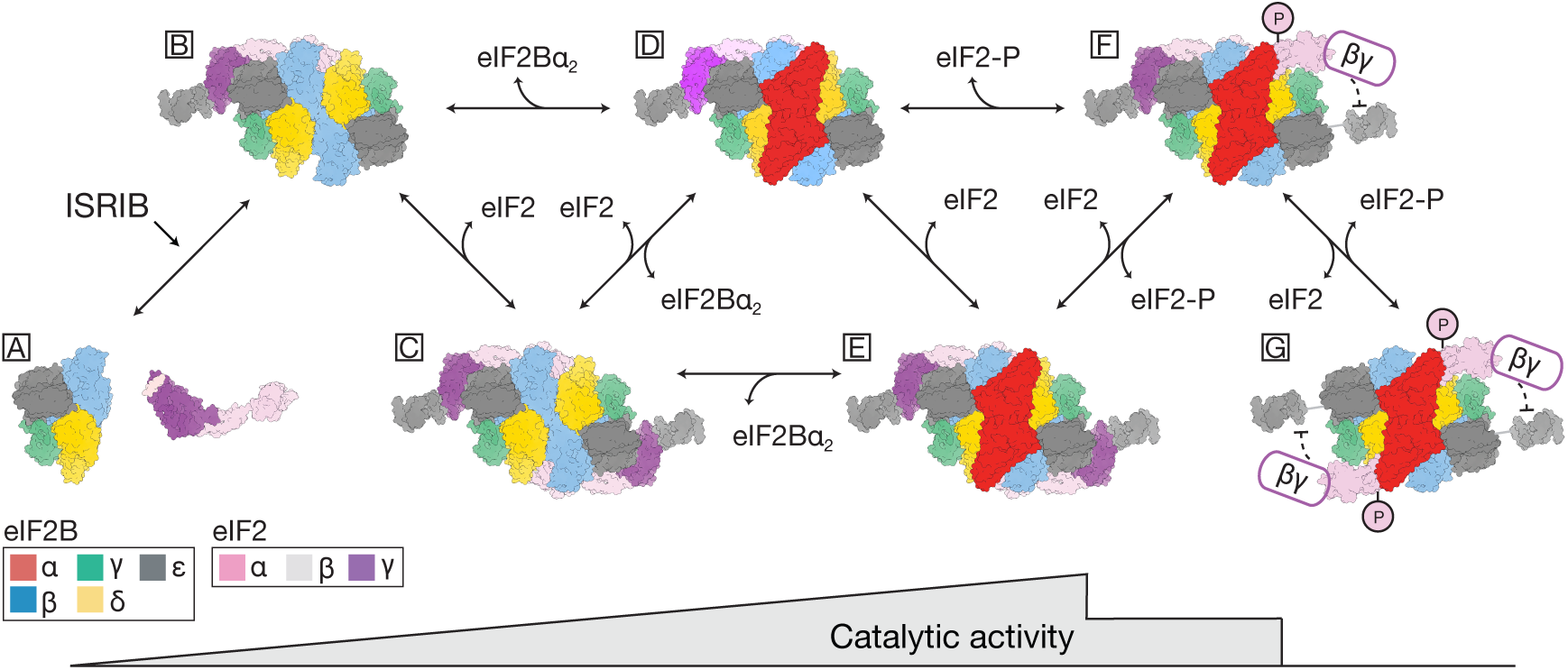
The structural basis of assembly-stimulated GEF activity and phosphoregulation by the ISR. Schematic representation of the regulated states of eIF2B assembly: dissociated tetramers and eIF2 heterotrimers (A), ISRIB and eIF2-stabilized octamers (B-C), fully assembled decamers (D-G), eIF2 substrate binding (B-F), and remodeling by phosphorylation (F,G). Relative catalytic activity of eIF2B-catalyzed nucleotide exchange upon oligomerization versus the stepwise loss of activity during inhibition by phosphorylated eIF2 (bottom).

These structures also deepen our understanding of ISRIB’s ability to ameliorate the inhibitory effects of eIF2α phosphorylation on ternary complex formation. ISRIB staples tetrameric building blocks together into an octamer, enhancing GEF activity three-fold, thus favoring association of the eIF2Bα_2_ homodimer. The summed effect of these sequential steps is a an order of magnitude activity enhancement (*20*). Thus, the surplus of GEF activity provided by ISRIB-driven holoenzyme assembly will counteract inhibition of eIF2B by limiting amounts of eIF2-P. By contrast, under conditions where eIF2B decamer is maximally stabilized at saturating concentrations of eIF2-P, ISRIB cannot promote any additional eIF2B decamer assembly. This mechanism explains ISRIB’s bell-shaped response profile to increasing stress (*19*), and its ability to mitigate the effects of eIF2 phosphorylation within a certain eIF2-P concentration range.

## Acknowledgments

Rights to the invention have been licensed by UCSF to Calico. We thank E. Pavolcak, J. Peschek, E. Karagöz, M. Boone, V. Belyy, G. Narlikar, R. Vale, the PHENIX and cryoSPARC developers, and members of the Walter and Frost labs for reagents, technical advice, and helpful discussions; M. Braunfeld, D. Bulkley, and M. Harrington of the UCSF Center for Advanced CryoEM, which are supported by NIH grants S10OD020054 and 1S10OD021741 and the Howard Hughes Medical Institute (HHMI); Z. Yu and H. Chou of the CryoEM Facility at the HHMI Janelia Research Campus; the QB3 shared cluster and NIH grant 1S10OD021596-01 for computational support; and G. Pavitt for the GP6452 yeast strain used in the purification of eIF2. A Titan X Pascal used for this research was donated by the NVIDIA Corporation.

## Funding

This work was supported by an HHMI Faculty Scholar grant (A.F.) and by Calico Life Sciences LLC, the Rogers Family Foundation, the Weill Foundation, and HHMI (P.W.). L.R.K. was supported by a graduate research fellowship from the N.S.F. A.F. is a Chan Zuckerberg Biohub Investigator, and P.W. is an Investigator of HHMI.

## Author contributions

Conception and design, analysis and interpretation of data: L.R.K., A.A.A., H.C.N., P.W. and A.F; acquisition of data: L.R.K., A.A.A., A.G.M.; writing (original draft): L.R.K., A.A.A., P.W., and A.F.; Writing (review and editing): L.R.K., A.A.A., H.C.N., A.G.M., C.J.K., L.A.M., J.C.T., L.E.M.V., A.F., and P.W.

## Competing interests

P.W. is an inventor on U.S. Patent 9708247 held by the Regents of the University of California that describes ISRIB and its analogs. Rights to the invention have been licensed by UCSF to Calico.

## Accession numbers

Accession numbers for the human eIF2•eIF2B structures are as follows: EMD-####, EMD-####, EMD-#### (density maps; Electron Microscopy Data Bank) and #### (coordinates of atomic models; Protein Data Bank).

## Data availability

All data needed to evaluate the conclusions in the paper are present in the paper and/or the supplementary materials, and the structural data is available in public databases. All of the raw cryo-EM particle images are available upon request. The GP6452 yeast strain is available under a material transfer agreement with the University of Manchester.

## Supplementary Materials

Materials and Methods Figures S1-S3

Tables S1-S3

Movie S1

References (*38-46*)

## Materials and Methods

### Purification of decameric eIF2B(αβγδε)2

As previously described (*20*), pJT066, pJT073, and pJT074 were co-transformed into One Shot BL21 Star (DE3) chemically competent E. coli cells (Invitrogen) and grown in Luria broth containing ampicillin, kanamycin, and chloramphenicol at 37°C on an orbital shaker. When the culture reached an OD600 of ~0.6, the temperature was reduced to 16°C, and the culture was induced with 0.8 mM IPTG (Gold Biotechnology) and grown for 16 hours. Cells were harvested and lysed with EmulsiFlex-C3 (Avestin) in a buffer containing 20 mM HEPES-KOH, pH 7.5, 250 mM KCl, 1 mM tris(2-carboxyethyl)phosphine (TCEP), 5 mM MgCl_2_, 15 mM imidazole, and complete EDTA-free protease inhibitor cocktail (Roche). The lysate was clarified at 30,000g for 20 min at 4°C. Subsequent purification steps were conducted on the ÄKTA Pure (GE Healthcare) system at 4°C. The clarified lysate was loaded onto a HisTrap HP 5 ml, washed in binding buffer (20 mM HEPES-KOH, pH 7.5, 200 mM KCl, 1 mM TCEP, 5 mM MgCl_2_, and 15 mM imidazole), and eluted with a linear gradient (75 ml) of 15 mM to 300 mM imidazole in the same buffer. The eIF2B fraction eluted from the HisTrap column at 80 mM imidazole. The eIF2B fraction was collected and loaded onto a 20 ml Mono Q HR16/10 column (GE Healthcare), washed in Buffer A (20 mM HEPES-KOH, pH 7.5, 200 mM KCl, 1 mM TCEP, and 5 mM MgCl_2_) and eluted with a linear gradient (200 ml) of 200 mM to 500 mM KCl in the same buffer. The eIF2B fraction eluted off the Mono Q column at a conductivity of 46 mS/cm (corresponding to 390 mM KCl). Fractions were collected, concentrated with an Amicon Ultra-15 concentrator (EMD Millipore) with a 100,000-dalton molecular weight cutoff, and loaded onto a Superdex 200 10/300 GL column (GE Healthcare) equilibrated with Buffer A. A typical preparation yielded approximately 0.5 mg of eIF2B(aβγδε)_2_ from a 1-liter culture.

### Purification of heterotrimeric human eIF2

Human eIF2 was prepared from an established recombinant *S. cerevisiae* expression protocol (*22*). In brief, the yeast strain GP6452 (gift from the Pavitt lab, University of Manchester) containing yeast expression plasmids for human eIF2 subunits and a deletion of GCN2 encoding the only eIF2 kinase in yeast, was grown to saturation in synthetic complete media (Sunrise Science Products) with auxotrophic markers (-Trp, -Leu, -Ura) in 2% dextrose. The β and α subunits of eIF2 were tagged with His6 and FLAG epitopes, respectively. A 12-liter yeast culture was grown in rich expression media containing yeast extract, peptone, 2% galactose, and 0.2% dextrose. Cells were harvested and resuspended in lysis buffer [100 mM Tris, pH 8.5, 300 mM KCl, 5 mM MgCl_2_, 0.1% NP-40, 5 mM imidazole, 10% glycerol (Thermo Fisher Scientific), 2 mM DTT, 1× protease inhibitor cocktail (Sigma Aldrich #11836170001), 1 µg/ml each aprotinin (Sigma Aldrich), leupeptin (Sigma Aldrich), pepstatin A (Sigma Aldrich). Cells were lysed in liquid nitrogen using a steel blender. The lysate was centrifuged at 10,000g for 1 hour at 4°C. Subsequent purification steps were conducted on the ÄKTA Pure (GE Healthcare) system at 4°C. Lysate was applied to a 5-ml HisTrap Crude column (Thermo Fisher Scientific) equilibrated in buffer (100 mM HEPES, pH 7.5, 100 mM KCl, 5 mM MgCl_2_, 0.1% NP-40, 5% glycerol, 1 mM dithiothreitol, 0.5× protease inhibitor cocktail, 1 µg/ml each aprotinin, leupeptin, pepstatin A). eIF2 bound to the column, was washed with equilibration buffer and eluted using a 50 ml linear gradient of 5 mM to 500 mM imidazole. Eluted eIF2 was incubated with FLAG M2 magnetic affinity beads, washed with FLAG wash buffer (100 mM HEPES, pH 7.5, 100 mM

KCl, 5 mM MgCl_2_, 0.1% NP-40, 5% glycerol, 1 mM TCEP, 1× protease inhibitor cocktail, 1 µg/ml each aprotinin, leupeptin, pepstatin A) and eluted with FLAG elution buffer [identical to FLAG wash buffer but also containing 3× FLAG peptide (100 µg/ml, Sigma Aldrich)]. Concentration of purified protein was measured by BCA assay (Thermo Fisher Scientific # PI23225); protein was flash-frozen in liquid nitrogen and stored in elution buffer at –80° C. A typical preparation yielded 1 mg of eIF2 from a 12-liter culture.

### Purification of human eIF2α

Human eIF2 was *E. coli* codon-optimized, synthesized and cloned into a pUC57 vector by GenScript Inc. PCR-amplified dsDNA fragments containing the eIF2α sequence were cloned into a pET28a vector using an In-Fusion HD Cloning Kit (Takara Bio), resulting in the kanamycin-resistant expression plasmid, pAA007. pAA007 was co-transformed into One Shot BL21 Star (DE3) chemically competent E. coli cells (Invitrogen), along with the tetracycline-inducible, chloramphenicol-resistant plasmid, pG-Tf2, containing the chaperones groES, groEL, and tig (Takara Bio). Transformed cells were grown in Luria broth containing kanamycin and chloramphenicol at 37°C on an orbital shaker.

When the culture reached an OD600 of ~0.2, 1ng/mL tetracycline was added to induce expression of chaperones. At an OD600 of ~0.8, the temperature was reduced to 16°C, eIF2α expression was induced with 1 mM IPTG (Gold Biotechnology) and the culture was grown for 16 hours. Cells were harvested and lysed with EmulsiFlex-C3 (Avestin) in a buffer containing 100 mM HEPES-KOH, pH 7.5, 300 mM KCl, 2 mM dithiothreitol (DTT), 5 mM MgCl_2_, 5 mM imidazole, 10% glycerol, 0.1% NP-40, and complete EDTA-free protease inhibitor cocktail (Roche). The lysate was clarified at 10,000g for 60 min at 4°C. Subsequent purification steps were conducted on the ÄKTA Pure (GE Healthcare) system at 4°C.

The clarified lysate was loaded onto a 5-ml HisTrap FF Crude column (GE Healthcare), washed in a buffer containing 20 mM HEPES-KOH, pH 7.5, 100 mM KCl, 5% glycerol, 1 mM DTT, 5 mM MgCl_2_, 0.1% NP-40, and 20 mM imidazole, and eluted with 75-ml linear gradient of 20 to 500 mM imidazole. The eIF2α containing fractions were then collected and applied to a MonoS HR 10/10 (GE Healthcare) equilibrated in a buffer containing 20 mM HEPES-KOH, pH 7.5, 100 mM KCl, 1 mM DTT, 5% glycerol, and 5 mM MgCl_2_. The column was washed in the same buffer and eluted with a 75-mL linear gradient of 100 mM to 1 M KCl. eIF2α containing fractions were collected and concentrated with an Amicon Ultra-15 concentrator (EMD Millipore) with a 30,000-dalton molecular mass cutoff and chromatographed on a Superdex 75 10/300 GL (GE Healthcare) column equilibrated in a buffer containing 20 mM HEPES-KOH, pH 7.5, 100 mM KCl, 1 mM TCEP, 5 mM MgCl_2_, and 5% glycerol. A typical preparation yielded approximately 2 mg of eIF2α from a 1-liter culture.

## Purification of phosphorylated human eIF2α

eIF2α and PKR were expressed and purified as above, but with the following modifications: One Shot BL21 Star (DE3) *E. coli* were co-transformed with pAA007, pG-Tf2, a third plasmid expressing the kinase domain of PERK (PERK 4: PERKKD-pGEX4T-1, Addgene plasmid #21817 donated by Dr. David Ron) and a resistance marker towards ampicillin. Transformed bacteria were grown in Luria broth containing ampicillin, kanamycin, and chloramphenicol. For purification, 1x PhosSTOP (Roche) was added to the lysis and purification buffers.

Phosphorylation was confirmed by Phos-Tag SDS-PAGE (Wako) as described previously (*38*), and by Western blot with an eIF2α S51 phosphorylation-specific antibody (Cell Signaling, #9721).

### Purification of tetrameric eIF2B(βγδε)

Tetrameric eIF2B(βγδε) and tetrameric eIF2B(βγδε) mutant proteins were purified using the same protocol as described for the decamer with the exception that expression strains were cotransformed without the eIF2B a subunit expressing plasmid. A typical preparation yielded approximately 0.75 mg of eIF2B(βγδε) from a 1-liter culture.

> eIF2B(βγδε) tetramer with co-transformed plasmids: pJT073, pJT074
>
> βN132D eIF2B(βγδε) tetramer with co-transformed plasmids: pAA012, pJT074
>
> δR250A eIF2B(βγδε) tetramer with co-transformed plasmids: pAA013, pJT074
>
> δR250E eIF2B(βγδε) tetramer with co-transformed plasmids: pAA014, pJT074

### Cloning of mutant eIF2B expression plasmids

Mutant eIF2B constructs were generated by site-directed mutagenesis on pJT073 for β and δ, and pJT066 for α, using the primer indicated and its reverse complement.

> βN132D (pAA012): 5′-CCACTACGCTCAGCTGCAGTCTGACATCATCGAAGCTATCAACG-3′ δR250A (pAA013): 5′-
>
> CCCCGCCGAACGAAGAACTGTCTGCTGACCTGGTTAACAAACTGAAACCG-3′ δR250E (pAA014): 5′-CCCCGCCGAACGAAGAACTGTCTGAGGACCTGGTTAACAAACTGAAACCG-3′

### EM sample preparation and data collection for ISRIB-bound eIF2•eIF2B and eIF2α•eIF2B complexes

Decameric eIF2B(αβγδε)_2_ + eIF2(αβγ) + ISRIB: eIF2B(αβγδε)_2_ was diluted to 800 nM eIF2B, eIF2 to 2 µM, and a stock solution of 200 µM ISRIB in N-methyl-2-pyrrolidone (NMP) was added to a final ISRIB concentration of 2 µM in a final solution containing 20 mM HEPES-KOH, pH 7.5, 100 mM KCl, 1 mM TCEP, 5 mM MgCl_2_, 0.1% NMP, and incubated on ice for 10 min. An inter-amine bifunctional crosslinker (Pierce premium BS3, #PG82084) was then added at a concentration of 0.25mM, and the mixture was incubated on ice for 2 hours before quenching with 10mM Tris HCl.

Decameric eIF2B(αβγδε)_2_ + eIF2α(P): eIF2B(αβγδε)_2_ was diluted to 800 nM and eIF2α(P) to 2.4 µM in a final solution containing 20 mM HEPES-KOH, pH 7.5, 100 mM KCl, 1 mM TCEP, 5 mM MgCl_2_, 0.1% NMP, and incubated on ice for 10 min and cross-linked as described above.

Each sample was applied to Quantifoil R 1.2/1.3 200 or 400 Au mesh grids (Quantifoil, Germany). Quantifoil grids were used without glow discharging. Using a Vitrobot Mark IV at 4°C and 100% humidity, 3.5 µl of sample was applied to the grid, incubated for an additional 10s, then blotted with 0 mm offset for ~6 s and plunge-frozen in liquid ethane. Two data sets were collected. Both data sets were collected with on a 300 kV Titan Krios at UCSF using a K2 Summit detector operated in super-resolution mode; 3233 images for eIF2P•eIF2B and 3947 images for eIF2•eIF2B were collected at a magnification of 29,000× (0.41Å per super-resolution pixel, binned by a factor of 2 to 0.82Å for subsequent processing). Dose-fractionated stacks were collected according to the parameters in Table S1.

## Image analysis and 3D reconstruction

All dose-fractionated image stacks were corrected for motion artifacts, 2× binned in the Fourier domain, and dose-weighted using MotionCor2 (*39*), resulting in one dose-weighted and one unweighted integrated image per stack with pixel sizes of 0.822 Å. The parameters of the contrast transfer function (CTF) were estimated using GCTF-v1.06 (*40*) and the motion-corrected but unweighted images; automated particle picking was done using Gautomatch-v0.53 and averaged in 2D using Cryosparc v0.6.5 (*41*). For the 3D reconstruction, an ab initio reconstruction was done without symmetry, followed by homogeneous refinement. High-resolution homogeneous refinement was then performed in cryoSPARC, using dynamic masks and imposed C2 symmetry for eIF2P bound to eIF2B and 2 eIF2 bound to eIF2B, C1 symmetry was used for 1 eIF2 bound to eIF2B. All maps were low-pass filtered and sharpened in cryoSPARC. Molecular graphics and analyses were performed with the UCSF Chimera package. UCSF Chimera is developed by the Resource for Biocomputing, Visualization, and Informatics and supported by NIGMS P41-GM103311 (*42*). Accession numbers for the structures are as follows: EMD-XXXX, EMD-XXXX, EMD-XXXX (density maps; Electron Microscopy Data Bank).

### Atomic modeling and validation

For all models, previously determined structures of the human eIF2B complex [PDB: 6CAJ (*8*)], human eIF2 alpha [PDBs: 1Q8K and 1KL9], the C-terminal HEAT domain of eIF2B epsilon [PDB: 3JUI], and mammalian eIF2 gamma [PDB: 5K0Y] were used for initial atomic interpretation. The models were manually adjusted and rebuilt in *Coot* (*43*) and then refined in phenix.real_space_refine (*44*) using global minimization, morphing, secondary structure restraints, and local grid search. Then iterative cycles of manually rebuilding in *Coot* and phenix.real_space_refine, with previous strategies and additionally B-factor refinement, were performed. The final model statistics were tabulated using Molprobity (*45*) (Table S3). Map versus atomic model FSC plots were computed after masking using EMAN2 (*46*) calculated density maps from e2pdb2mrc.py with heteroatoms (ISRIB) and per-residue B-factor weighting. Solvent accessible surfaces and buried surface areas were calculated from the atomic models using UCSF Chimera. Final atomic models have been deposited at the PDB with the following accession codes: 1 eIF2•eIF2B•ISRIB (xxxx); 2 eIF2•eIF2B•ISRIB (xxxx); phosphorylated eIF2α•eIF2B (xxxx). All structural figures were generated with UCSF Chimera (*42*) and BLENDER (http://www.blender.org).

### GDP exchange assay

In vitro detection of GDP binding to eIF2 was adapted from a published protocol for a fluorescence intensity–based assay describing dissociation of eIF2 and nucleotide (*14*). We modified the procedure to establish a loading assay for fluorescent GDP as described (*20*). For the “GDP loading assay,” purified eIF2 (200 pmol) was incubated with a molar equivalent Bodipy-FL-GDP (Thermo Fisher Scientific) in assay buffer (20 mM HEPES, pH 7.5, 100 mM KCl, 5 mM MgCl_2_, 1 mM TCEP, 0.1% NMP, and 1 mg/ml bovine serum albumin) to a volume of 18 µl in 384 square-well black-walled, clear-bottom polystyrene assay plates (Corning). The reaction was initiated by addition of 2 µl of buffer or purified wild-type and mutant eIF2B(βγδε) (2 pmol) under various conditions to compare nucleotide exchange rates. For comparison of tetramer or ISRIB-assembled octamer, eIF2B(βγδε) (2 pmol) was preincubated in 0.1% NMP and 2 mM ISRIB for 15 min before 10-fold dilution into the final reaction. These concentrations of vehicle and ISRIB were used throughout unless otherwise specified. Fluorescence intensity for both loading and unloading assays was recorded every 10 s for 60 or 100 min using a TECAN M1000 Pro plate reader (excitation wavelength: 495 nm, bandwidth 5 nm, emission wavelength: 512 nm, bandwidth: 5 nm). Data collected were fit to a first-order exponential.

---

**Fig. S1.**
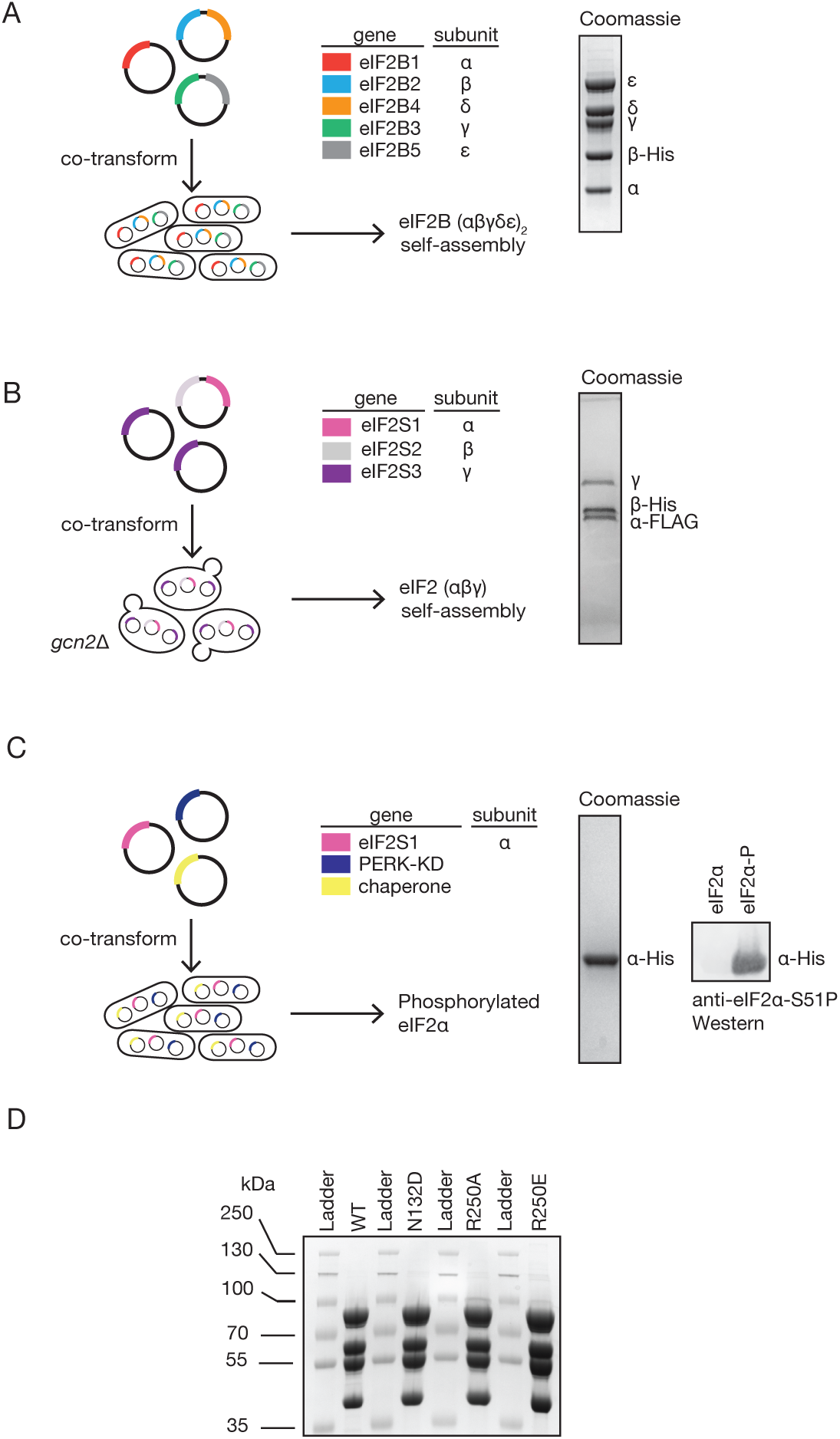
Purification of eIF2, eIF2α-P, eIF2B(αβγδε)_2_, and mutant eIF2B(βγδε) (A) Recombinant *E. coli* expression system and SDS-PAGE analysis for human eIF2B(αβγδε)_2_ as described in (*20*). (B) Recombinant *S. cerevisiae* expression system and SDS-PAGE analysis for human eIF2 as described in (*22*). (C) Recombinant expression *E. coli* system for phosphorylated eIF2α and SDS-PAGE and Western blot analysis as described in (*38*). This expression protocol was modified to include the chaperones GroEL, GroES and tig. (D) SDS-PAGE analysis of mutant eIF2B(βγδε).

**Fig. S2.**
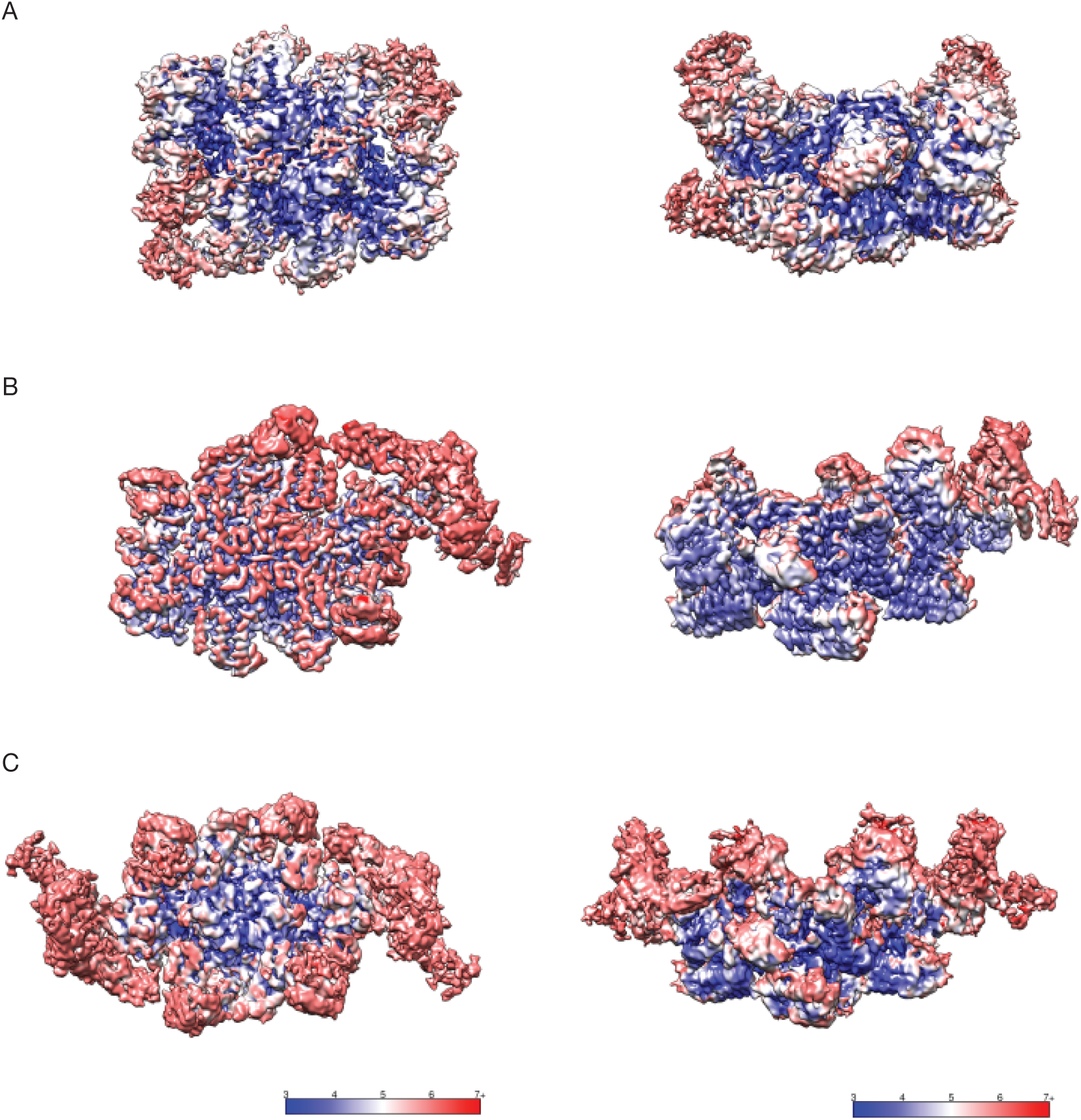
Local resolution Estimates. (A) Local resolution estimates determined using cryoSPARC v0.6.5 and displayed using UCSF Chimera for eIF2α-P bound to eIF2B with C2 symmetry, (B) eIF2 bound to eIF2B with C1 symmetry and (C) two copies of eIF2 bound to eIF2B with C2 symmetry.

**Fig. S3.**
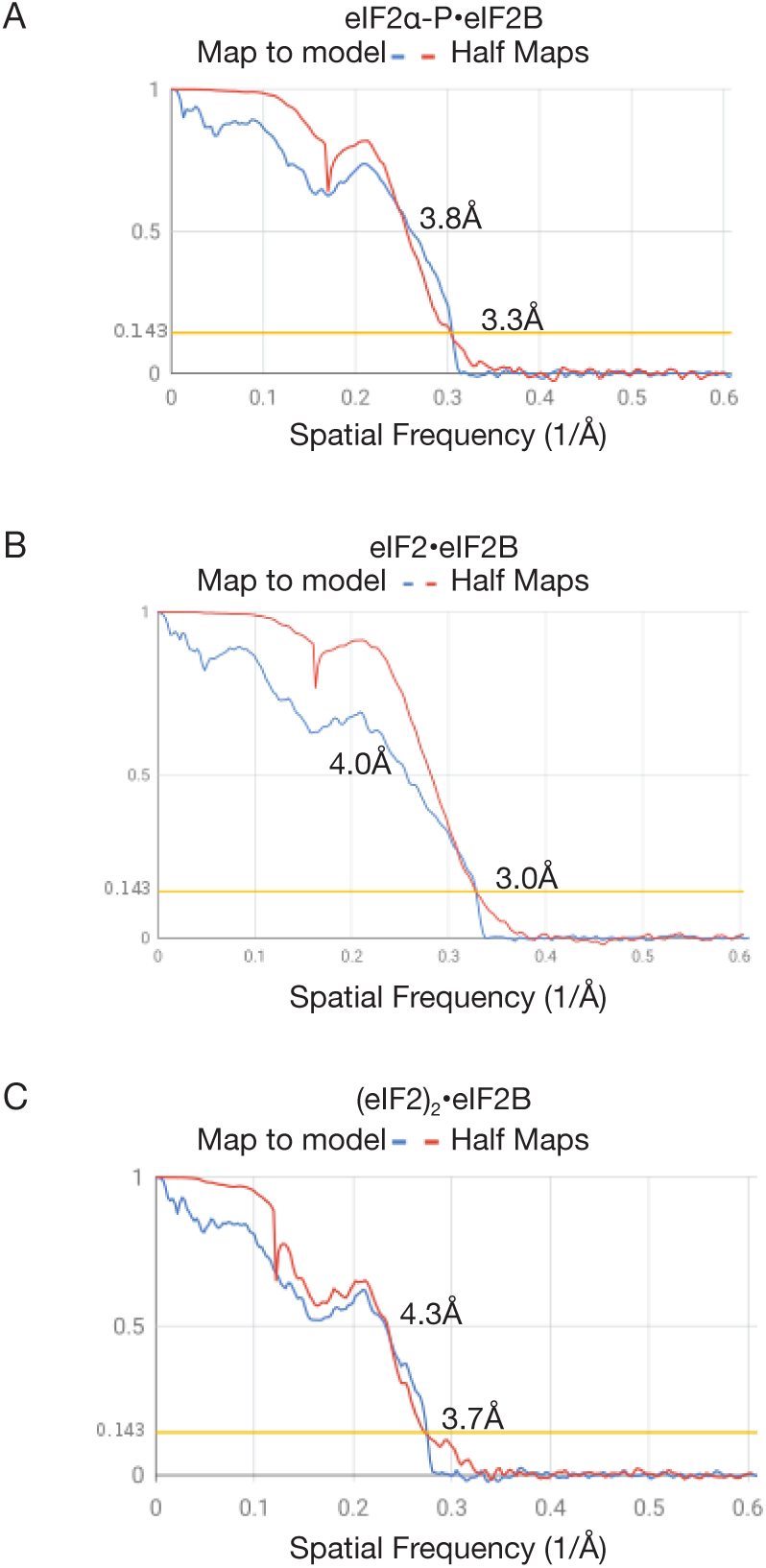
Fourier shell correlations. (A) eIF2α-P bound to eIF2B with C2 symmetry. Correlations between the independent half maps (red), and the final cryoEM density map versus simulated density maps for the atomic models (blue). (B) eIF2 bound to eIF2B with C1 symmetry, and (C) two copies of eIF2 bound to eIF2B with C2 symmetry.

**Table S1.** Data collection parameters.

|  | eIF2 Alpha P bound to eIF2B | eIF2 bound to eIF2B and Two eIF2 bound to eIF2B |
| --- | --- | --- |
| Pixel Size (Å) | 0.822 | 0.822 |
| Defocus Range | 0.5 - 1.5 | 0.5 - 1.5 |
| Voltage (kV) | 300 | 300 |
| Magnification | 29,000 | 29,000 |
| Spherical Aberration (mm) | 2.7 | 2.7 |
| Detector | K2 Summit | K2 Summit |
| Detector Pixel Size | 5 | 5 |
| Per frame electron dose (e-/Å<sup>2</sup>) | 0.8 | 0.85 |
| # of Frames | 100 | 80 |
| Frame Length (seconds) | 0.1 | 0.1 |
| Micrographs | 3233 | 3947 |
| Exposure (seconds) | 10 | 8 |

**Table S2.** Refinement parameters.

|  | eIF2 Alpha P bound to eIF2B | eIF2 bound to eIF2B | Two eIF2 bound to eIF2B |
| --- | --- | --- | --- |
| Particles | 22165 | 44157 | 7,473 |
| FSC Average Resolution, unmasked (Å) | 3.48 | 3.15 | 3.78 |
| FSC Average Resolution, masked (Å) | 3.27 | 3.04 | 3.65 |
| Map Sharpening B-factor | -78.7 | -71.3 | -41.8 |

**Table S3.** Modeling statistics.

|  | eIF2 Alpha P bound to eIF2B | eIF2 bound to eIF2B | Two eIF2 bound to eIF2B |
| --- | --- | --- | --- |
| Number of Atoms, macromolecules | 25100 | 28495 | 33629 |
| Number of Atoms, ligands | 0 | 60 | 60 |
| Molprobity Score | 2.2 | 1.28 | 1.53 |
| Clashscore, all atoms | 6.61 | 1.86 | 4.27 |
| Outlier Rotamers (%) | 3.16 | 0.12 | 0.26 |
| RMS (bonds) | 0.012 | 0.003 | 0.003 |
| RMS (angles) | 1.215 | 0.804 | 0.742 |
| Ramachandran Favored (%) | 92.82 | 95.24 | 95.39 |
| Ramachandran Outliers (%) | 0 | 0 | 0 |
| Ramachandran Allowed (%) | 7.18 | 4.76 | 4.61 |

## Movie S1

Conformational morphing between the non-phosphorylated and S51-phosphorylated structures of eIF2α, highlighting how phosphorylation leads to refolding of the S-loop and the exposure of a hydrophobic surface.

